# Convergent genome evolution reveals shared routes to salt tolerance in flowering plants

**DOI:** 10.64898/2026.10.12.758530

**Authors:** Samadhi Wimalagunasekara, Maheshi Dassanayake

## Abstract

How do plants repeatedly evolve the ability to thrive in saline environments? We analyzed ∼10 million protein-coding genes from 232 angiosperm genomes spanning 55 families to identify convergent genomic signatures of halophytism. Across lineages, halophytes exhibit recurrent patterns in salt-response genes, including reduced gene numbers, higher basal expression, and constrained expression variation. We identified distinct evolutionary rate patterns and independent positive selection at the same amino acid sites across halophytes, revealing molecular convergence. Comparisons of crops and their wild halophytic relatives uncover contrasting gene family trajectories in stress-response pathways. These findings challenge the idea that increased gene number is the predominant route to adaptation and reveal recurrent genomic strategies associated with halophytism, advancing understanding of convergent evolution and providing a foundation for developing salt-tolerant crops.

## Introduction

Halophytism has evolved repeatedly across angiosperms, emerging independently in multiple lineages over different evolutionary timescales (*1*). Halophytes occupy ecologically diverse saline habitats, from coastal wetlands to saline deserts, and span growth forms from herbs to trees. This ecological and morphological diversity suggests that salt tolerance can arise through multiple evolutionary routes (*2–4*). Yet, salt tolerance remains one of the most demanding challenges for plant life, requiring coordinated regulation of energy cost between salt stress responses and growth that vary among species (*5–7*). Despite these challenges, halophytism occurs across numerous plant families, with halophytes estimated to comprise approximately 0.25% of all plant species (*8*). This repeated evolution across independent lineages provides a powerful natural framework to investigate different genomic solutions for salt tolerance.

Across halophytes, recurrent exposure to saline environments has favored similar structural and physiological adaptations despite their independent evolutionary origins (*8*). However, phenotypic convergence can arise through repeated recruitment of the same gene families across lineages or through lineage-specific changes in distinct genes and regulatory networks (*9–11*). Whether halophytes share a common genomic toolkit for salt tolerance or have evolved distinct molecular solutions, or a combination of both, remains poorly understood. Comparative genomic analyses across diverse lineages therefore provide an opportunity to identify recurrent evolutionary patterns and establish the extent to which salt adaptation reflects convergence at the genomic level.

Here, we curated a broad collection of angiosperm genomes spanning over 20 plant orders to identify genomic features associated with halophytism relative to non-halophytic relatives. We assessed convergence across three genomic dimensions: gene copy number allocation to salt-associated functions, basal expression of stress-response genes, and evolutionary rates and signatures of positive selection. This integrative approach provides a comparative framework for distinguishing recurrent genomic strategies from lineage-specific evolutionary changes, advancing our understanding of the genomic basis of convergent salt adaptation across angiosperms.

## Result

### A comparative catalog of ∼10 million genes across 232 angiosperm genomes

To establish a systematic comparative resource for testing whether angiosperm gene composition bears signatures of convergent evolution, we systematically selected 232 high-quality genomes spanning 55 angiosperm families, including halophytes, salt-sensitive relatives, and two basal salt-sensitive outgroups (*Nymphaea colorata* and *Amborella trichopoda*) (Fig. 1A). Halophytes were defined as plants capable of growing at ≥200 mM NaCl (*1*). We applied genome-wide gene model prediction (hereafter, genes) and functional annotation to construct a curated angiosperm gene catalog. The genome selection and catalog construction workflow is detailed in the Materials and Methods (Fig. S1, Table S1). The resulting gene catalog comprises ∼10 million genes, including 4.77 million from halophytes and 4.81 million from non-halophytes.

**Fig. 1.**
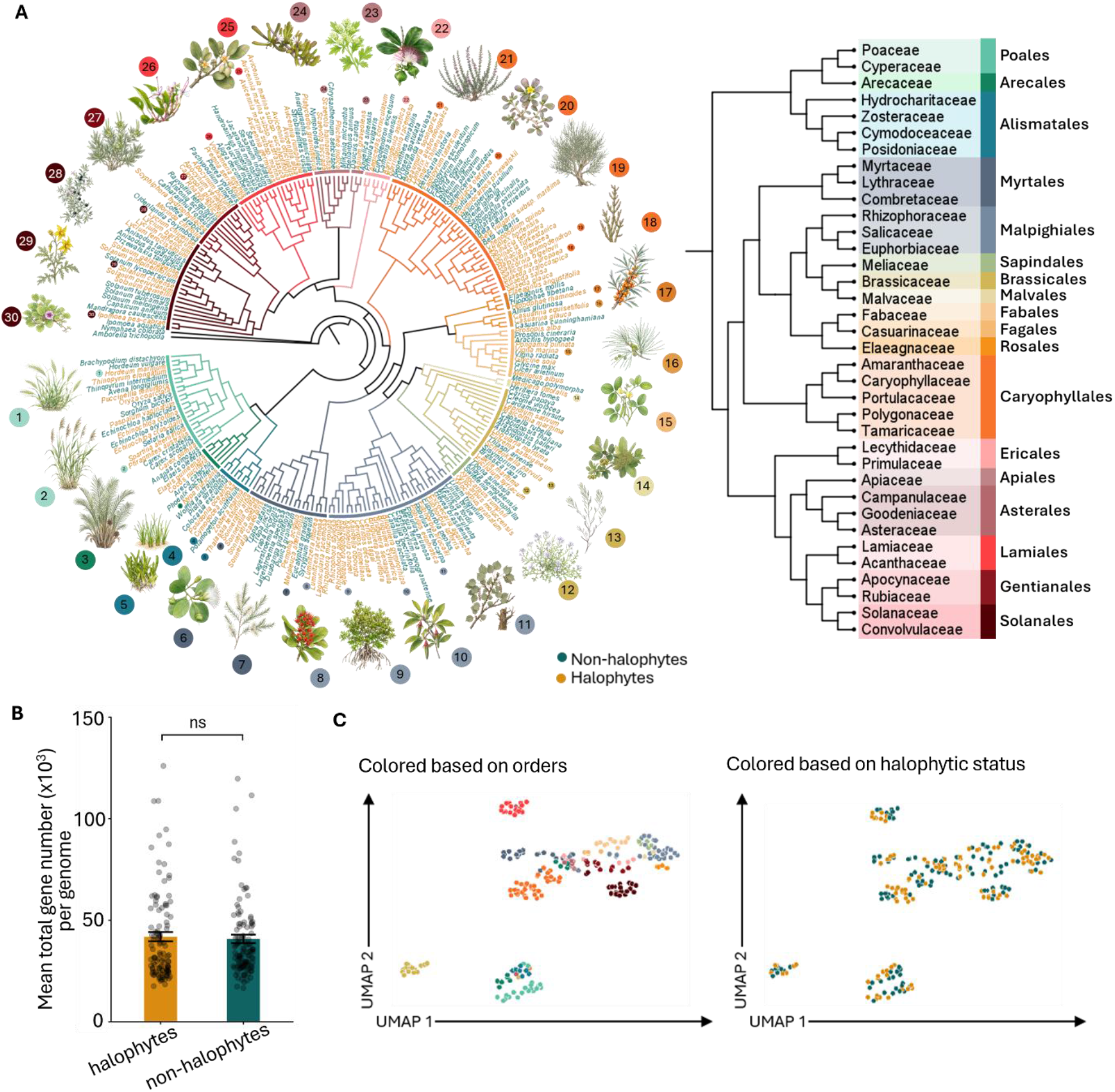
A comparative angiosperm gene catalog spanning 232 genomes in 55 families. **(A)** Phylogenetic relationships among the angiosperm genomes representing ∼10 million protein-coding genes from 114 halophytes (gold) and 118 non-halophytes (teal) with representative halophyte illustrations displayed around the phylogeny. Branches colored based on 18 plant orders listed in the tree on the right. **(B)** The total number of genes per genome among halophytes and non-halophytes. Significant differences tested at FDR-adjusted P < 0.05 based on Wilcoxon rank-sum tests with Benjamini–Hochberg correction; ns, not significant. Error bars indicate mean ± SD. **(C)** Clustering of species based on the number of genes in orthogroups represented in >25% of species. The upper and lower panels are colored by plant order and halophytic status, respectively, showing that species cluster primarily according to phylogenetic relationships.

The mean number of genes per genome did not differ between halophytes and non-halophytes (Fig. 1B), reducing the likelihood that gross annotation differences drive clustering. Gene copies assigned to 327,031 orthogroups (Data S1 and S2) were distributed differently among ortholog classes across lineages (Fig. S2, A and B; Table S2), with notable differences between halophytes and non-halophytes in the proportions of species-specific paralogs and single-copy orthologs (Fig. S2B). Species clustered primarily by phylogeny when restricting to orthogroups present in ≥25–50% of species (Fig. 1C and Fig. S2C).

### Less is more: reduced salt response gene representation in halophytes

Angiosperms have diversified over at least 150 Mya, adapting to diverse geographic, climatic, and abiotic conditions (*12*). Functional studies in model plants have established roles for thousands of genes in salt responses (*13*). We therefore selected ten Gene Ontology (GO) functional groups closely associated with salt responses and applied a standardized annotation pipeline to identify candidate salt-response genes in our catalog (Fig. 2A and Table S3). This yielded 338,577 genes, representing 3.3% of the catalog. We then assessed whether principal components (PCs) derived from these genes could distinguish halophytes from non-halophytes. We used binomial regression to identify the best-performing PC pair and evaluated classification accuracy using the area under the receiver operating characteristic curve. This analysis distinguished halophytes from non-halophytes despite substantial phylogenetic variation across angiosperms (Fig. 2B).

**Fig. 2.**
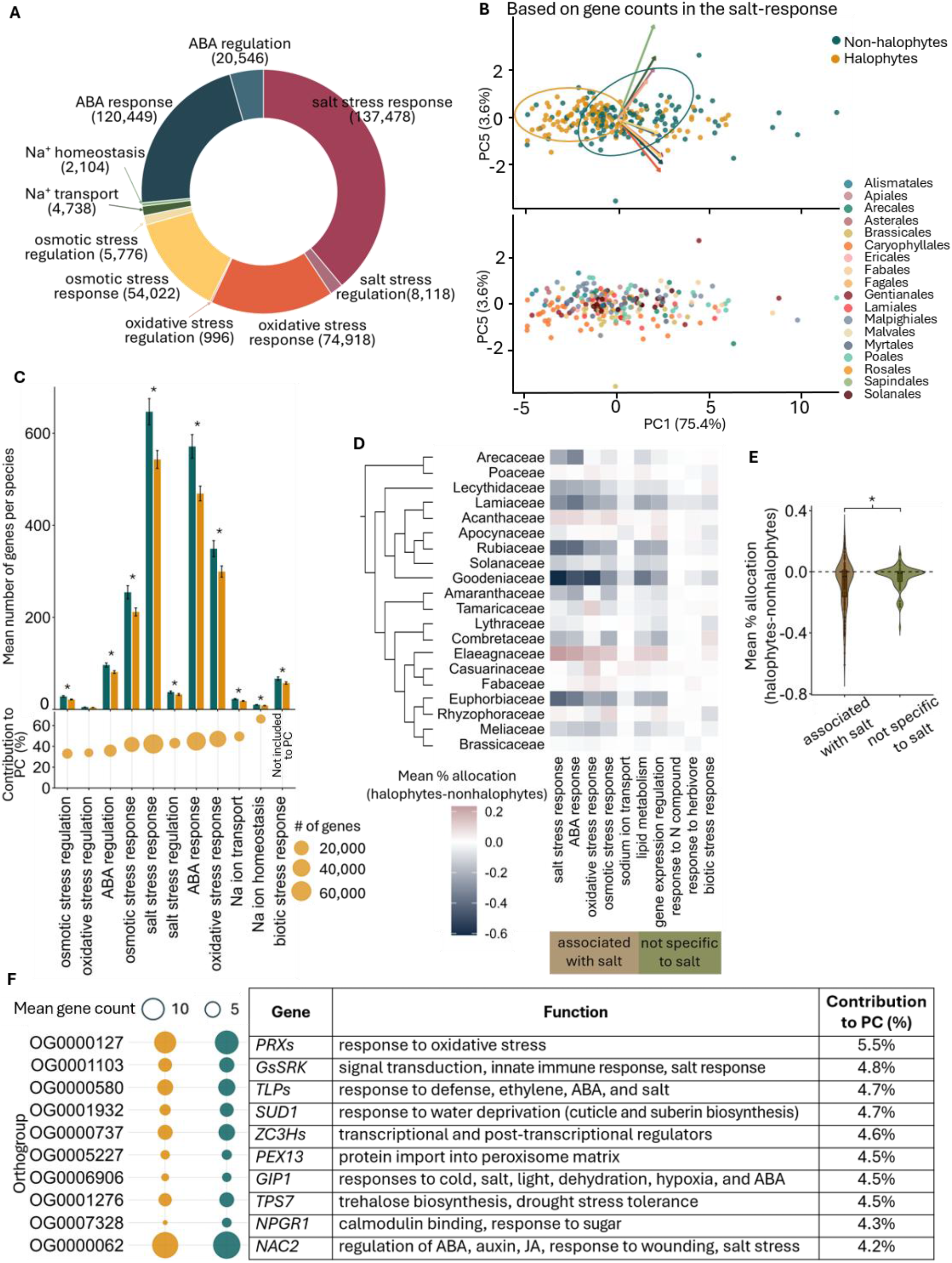
Reduced gene copy numbers characterize halophytes, particularly in salt-response functions. **(A)** Gene Ontology (GO) functional categories associated with salt responses and the numbers of genes assigned to each category across the 232 species. **(B)** Principal components (PCs) most influential in separating halophytes from non-halophytes based on gene counts in the salt-response categories shown in (A). Cohen’s *d* and Benjamini–Hochberg false discovery rate (BH-FDR)-adjusted *P* values (*p*adj) were used to quantify the separation between halophytes and non-halophytes along individual PCs. Logistic regression identified PC1 and PC5 as the pair with the highest area under the receiver operating characteristic curve (AUC). Species are colored by halophytic status (upper panel) or plant order (lower panel). Arrows indicate the contributions of individual functional categories to the separation, with arrow colors corresponding to the categories in (A). **(C)** Mean number of genes per species assigned to salt-response functions and biotic stress-response functions, the latter serving as a comparative category not specific to salt responses. Bubble sizes indicate the relative contributions of these functions to the PCs in (B). Asterisks indicate significant differences between halophytes and non-halophytes based on Wilcoxon rank-sum tests with BH-FDR-adjusted *P*adj < 0.05. Error bars indicate mean ± SD. **(D)** Heatmap of differences in mean percentage gene allocation between halophytes and non-halophytes for five selected salt-response functions and five other biological functions not specifically associated with salt responses. Positive values (pink) indicate greater relative gene allocation in halophytes, whereas negative values (blue) indicate greater relative allocation in non-halophytes. **(E)** Distribution of differences in mean percentage gene allocation between halophytes and non-halophytes for salt-response functions and functions not specifically associated with salt responses used in (D). Significance was assessed using Student’s *t*-test (*P* < 0.05). **(F)** Ten orthogroups contributing most to the separation of halophytes and non-halophytes in (B), with orthogroup IDs, functional annotations, and percentage contributions to the PCs. Bubble sizes indicate the mean gene count per orthogroup.

Genes associated with Na⁺ homeostasis and transport contributed most to distinguishing halophytes from non-halophytes, at 66.31% and 49.52%, respectively (Fig. 2C). Contrary to our expectation that halophytes would have higher gene copy numbers in key salt-response processes, they had fewer genes in all selected functional categories except oxidative stress regulation (Fig. 2C). We tested whether this reduction reflected a broader pattern across other biological processes, as suggested by the lower allocation of genes to biotic stress-response genes (Fig. 2C) or whether it was more pronounced for salt-associated functions. We also tested whether the reduced gene copy allocation to salt-response functions reflected an angiosperm-wide pattern among halophytes or was driven by a few families. To assess this, we examined 20 angiosperm families represented by more than four genomes, each with approximately equal representation of halophytes and non-halophytes. We compared equal numbers of salt-associated functional categories and biological processes not specifically associated with salt responses (see Methods). The results were consistent with the overall pattern of fewer gene copies allocated to salt-response functions in halophytes, although exceptions occurred in families such as Acanthaceae and Elaeagnaceae (Fig. 2, D and E). The recurrence of this pattern across multiple families suggests that reduced gene copy allocation to salt-response functions may represent a convergent genomic trend among halophytes.

Having established that gene composition distinguishes halophytes from non-halophytes across phylogenetic boundaries, we next sought to identify the orthogroups driving this separation without biasing toward genes already annotated for salt responses. To identify convergent signatures for halophytism beyond phylogenetic variation, we prioritized core, conserved genes represented by many species over lineage-specific, ecology-linked gene families that are also often poorly characterized functionally. We reasoned that halophytes might share orthogroups absent from Arabidopsis or rice, the primary sources of functional gene annotations, and thus overlooked by existing Gene Ontology annotations. Conversely, genes assigned to other functions in salt-sensitive ancestors may have been independently co-opted for salt responses in halophytes. We therefore examined orthogroups present in at least 50% of sampled species and ranked them by their contributions to discriminating halophytes from non-halophytes (Fig. S3). Of the 8,325 orthogroups represented in at least 50% of species, 7,570 (∼90%) had fewer gene copies contributed by halophytes than by non-halophytes, indicating a widespread trend in gene representation. The top 648 orthogroups (7.7% of those examined) accounted for 18.31% of the cumulative discriminatory contribution, marking an inflection point in the contribution profile (Table S4). We designate these orthogroups as “keystone” gene families because their gene composition contributes disproportionately to distinguishing halophytes from non-halophytes. Several of these lack functional annotations, highlighting potentially overlooked gene families that are broadly represented across angiosperms that are retained in halophytes. Among the annotated keystone families, the ten highest-ranking orthogroups were associated with broad stress-tolerance functions (Fig. 2F), paralleling the role of keystone gene families in contributing to multiple metabolic traits in yeast (*14*).

### High but stable: convergent elevation of basal expression in halophyte salt-response genes

We processed 239 publicly available RNA-seq datasets representing shoot tissues of 107 halophytes and 105 non-halophytes sampled under non-stress conditions to estimate basal expression (see Methods). Transcriptome-wide, mean gene expression declined with increasing gene number per genome in both groups, but the decline was steeper in halophytes than in non-halophytes (Fig. 3A and Table S5). Halophytes showed a lower coefficient of variation (CV) than non-halophytes, indicating more stable transcriptome-wide expression variability (Fig. 3B and Table S6). This pattern was maintained in CV of mean expression per species for the majority of salt-related functional categories (Fig. 2A, 3C and Table S7).

**Fig. 3.**
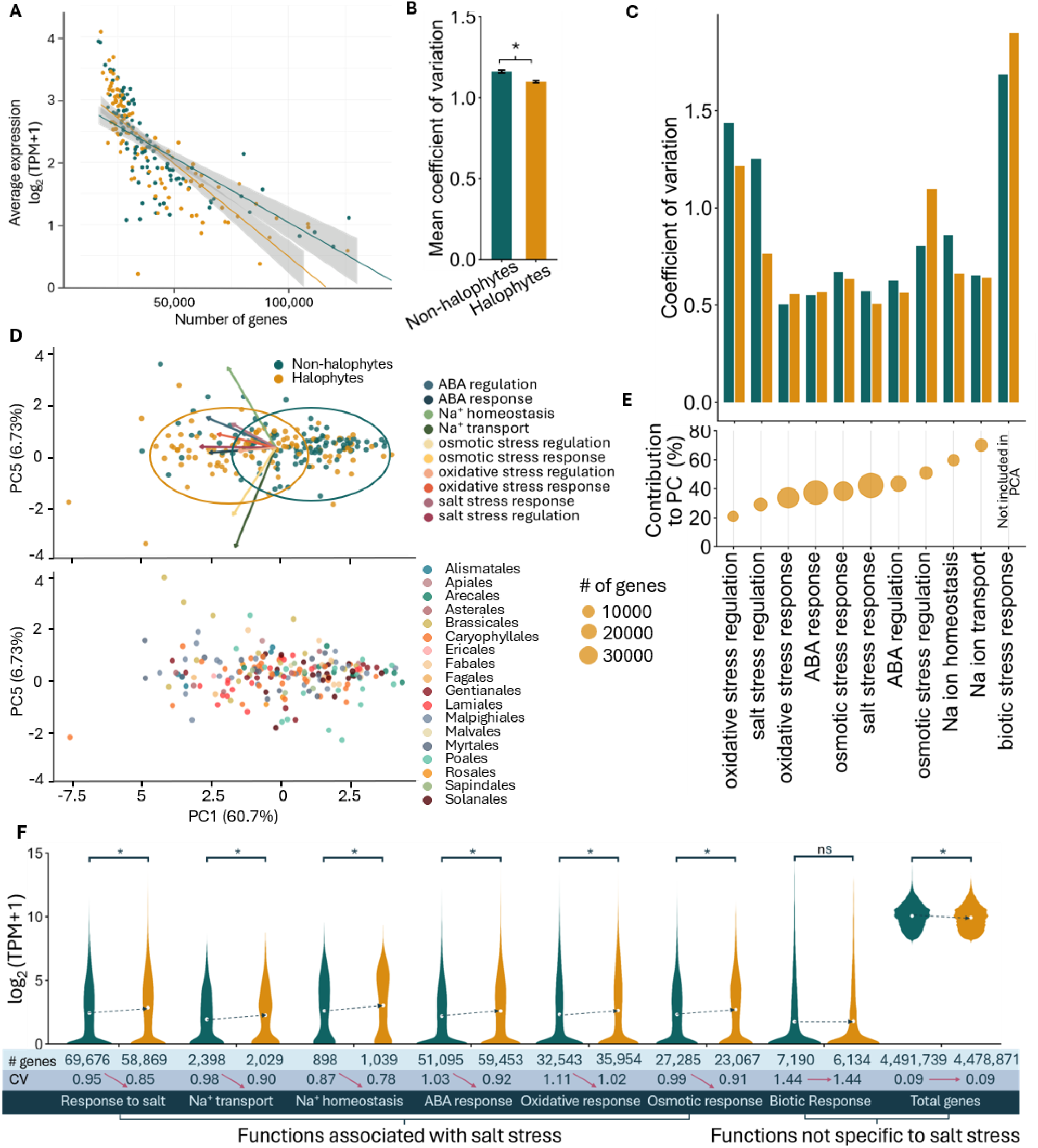
Convergent elevation and constraint of basal expression in halophytes compared to non-halophytes. **(A)** Transcriptome-wide mean expression based on the total number of genes in each genome. Points represent individual species. The solid lines indicate linear regression, and shaded regions represent 95% confidence intervals. **(B)** Mean coefficient of variation in basal expression of halophytes and non-halophytes. Asterisk indicates a significant difference based on the Wilcoxon rank-sum test (p < 0.05). Error bars show mean ± SD. **(C)** Coefficient of variation of basal expression for salt-response genes and biotic stress-response functions, the latter serving as a comparative category not specific to salt responses. **(D)** Principal components (PCs) most influential in separating halophytes from non-halophytes based on basal expression associated with salt-response genes. Cohen’s d and Benjamini–Hochberg false discovery rate (BH-FDR)-adjusted P values (*p*adj) were used to quantify the separation between halophytes and non-halophytes along individual PCs. Logistic regression identified PC1 and PC5 as the pair with the highest area under the receiver operating characteristic curve (AUC). Species are colored by halophytic status (upper panel) or plant order (lower panel). Arrows indicate the contributions of individual functional categories to the separation. **(E)** Bubble sizes indicate the relative contributions of these functions to the PCs in (D). **(F)** Basal expression distributions of genes assigned to salt-response functions and biotic response for halophytes (gold) and non-halophytes (green). CV-Coefficient of variation. Dashed arrows connect the mean basal expression and indicate the directional differences, and red arrows show the change of direction in CV. Significant differences between halophytes and non-halophytes were tested using the Wilcoxon rank-sum test. Asterisks indicate Benjamini–Hochberg FDR-adjusted P-values < 0.05. ns: not significant.

We next tested whether basal expression of salt-response genes could distinguish halophytes from non-halophytes. PC-based analyses discriminated the two groups despite phylogenetic variation, with selected PCs providing the greatest separation (Fig. 3D and Table S8). Strikingly, the functional categories contributing most to this separation mirrored those identified from gene copy number (Fig. 2C): Na⁺ transport contributed most (70%), followed by Na⁺ homeostasis (59%) (Fig. 3E and Table S8). Halophytes also exhibited higher mean basal expression and lower coefficients of variation for salt-response genes than non-halophytes (Fig. 3F and Table S9). This pattern was specific to salt-response genes and was not observed for genes associated with other functions, including biotic responses, or for global mean expression. Thus, halophytes show higher and more stable basal expression of selected functions associated with salt tolerance and not uniformly elevated overall expression, indicating a convergent pattern.

To assess whether basal expression could also distinguish halophytes from non-halophytes independently of existing functional annotations, we analyzed expression across all orthogroups represented in at least 50% of sampled species (Fig. S4A and Table S10). For orthogroups containing multiple genes from a species, we used the highest expression value among those genes to represent that species (Data S3 and S4). The orthogroups based on expression contributing most to the separation of halophytes from non-halophytes were primarily associated with general stress responses and energy metabolism (Fig. S4B). The general stress-response orthogroups mirrored the functions of gene families that contributed to the separation of the two groups based on gene numbers (Fig. 2F). Overall, 7,790 orthogroups (2.38% of all orthogroups, which includes 37.2% out of 10 million genes) showed convergent patterns among halophytes by lower gene copy number contributions, higher basal expression, or both in salt-response functions.

### Slow and specific: convergent evolutionary constraint, recurrent positive selection, and functional convergence in halophytes

Although previous studies have reported accelerated evolution and positive selection in ion transporters associated with salt tolerance, it remains unclear whether these patterns reflect lineage-specific adaptations or recur broadly across angiosperms (*15*, *16*). In our study, Na⁺ transport and Na⁺ homeostasis were among the most influential functions distinguishing halophytes from non-halophytes, motivating us to test whether salt-response genes exhibit convergent shifts in relative evolutionary rates shared by halophytes. We focused on Brassicaceae and Poaceae, representing eudicots and monocots, respectively. They contained the largest numbers of paired halophyte and non-halophyte genomes in our dataset, providing a comparative framework to assess evolutionary patterns associated with halophytism. We analyzed orthogroups represented in at least 50% of sampled species within Brassicaceae (17,402 orthogroups; Data S5 and S6) and Poaceae (16,327 orthogroups; Data S7 and S8) to identify relative evolutionary rate differences between halophytes and non-halophytes (see Methods). When examining evolutionary rates applied to the entire gene length, we identified 2,556 Brassicaceae and 3,102 Poaceae orthogroups in which genes from halophytes exhibited different evolutionary rates than those from their non-halophyte relatives (Fig. 4A; Tables S11 and S12). We identified inflection points in the distribution of relative evolutionary rates in halophytes compared with non-halophytes to classify orthogroups as fast- or slow-evolving (Fig. S5; Tables S11 and S12). The slow-evolving set contained more genes than the fast-evolving set in both Brassicaceae and Poaceae. Functional enrichment analyses further revealed that salt-response genes were more strongly enriched among slow-evolving than fast-evolving genes in both families (Fig. 4B and Table S13). This shared bias was evident for genes associated with abscisic acid (ABA) pathway regulation, antioxidant functions, and Na⁺/K⁺ homeostasis. Despite strong convergence in functional categories, the specific genes recruited and the molecular mechanisms underlying these shared functions varied substantially across lineages, indicating that similar salt-adaptation outcomes may emerge through diverse evolutionary routes.

**Fig. 4.**
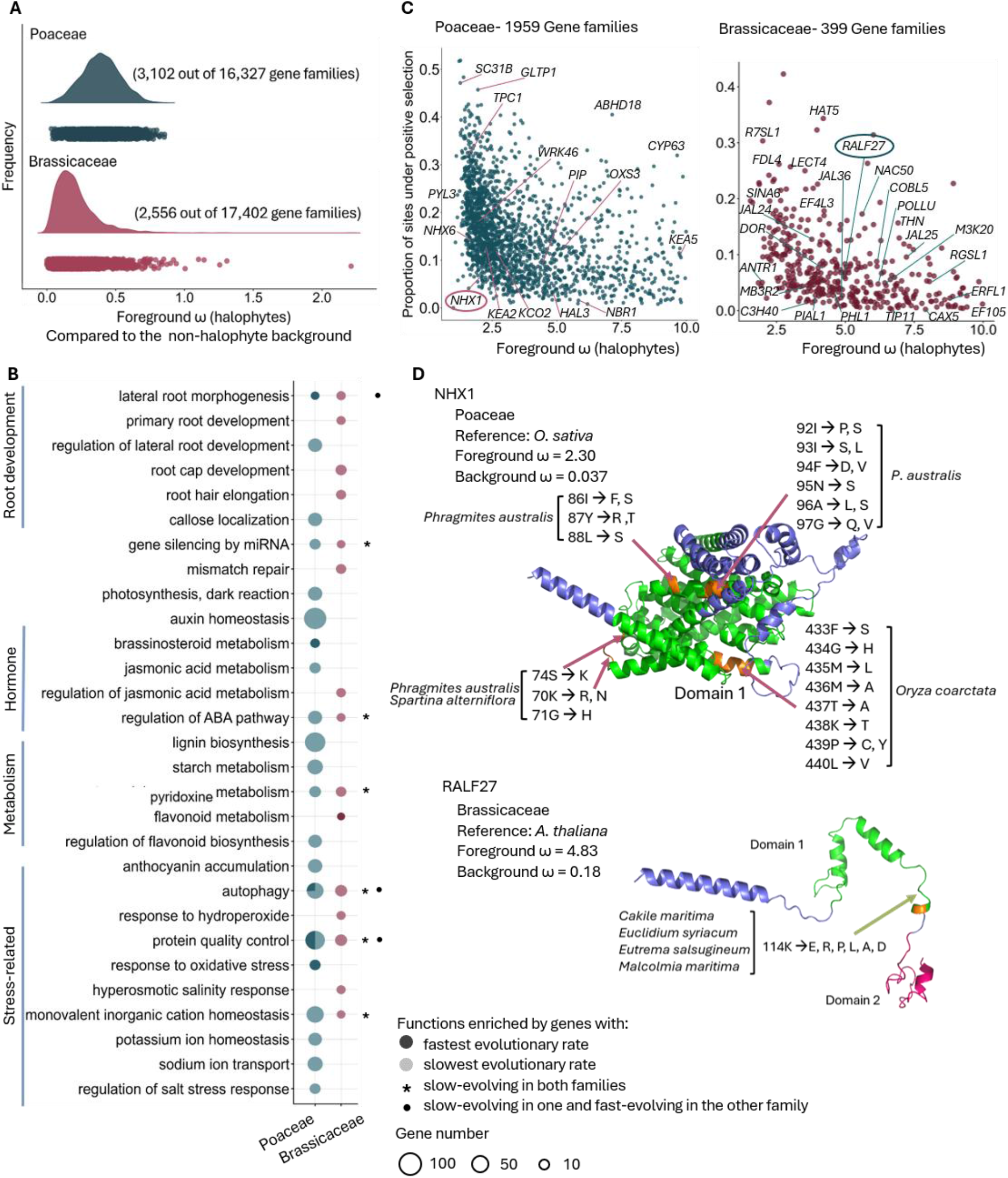
Convergent evolutionary rates and sites selected in halophyte genes associated with salt responses in Brassicaceae and Poaceae. **(A)** Distribution of halophyte ω values estimated from branch-model analyses for orthogroups in Poaceae (blue) and Brassicaceae (red). (B) Functional enrichment of genes with the fastest and slowest evolutionary rates in Poaceae and Brassicaceae halophytes. * indicates the functions enriched by the slow-evolving genes in both families, and • indicates functions enriched by slow-evolving genes in one family and fast-evolving genes in the other family. The bubble size indicates the number of genes. Enriched functions were identified with significance assessed using FDR-adjusted P-values < 0.05. **(C)** Positively selected genes estimated from the branch-site model in halophytes compared to non-halophytes in Poaceae (blue) and Brassicaceae (red). Orthogroups that are fitted to the alternative model after the Chi-square testing and FDR correction (*p*adj < 0.05) and 10> ω > 1 are plotted. **(D)** Positively selected sites in well-characterized salt-associated genes mapped onto predicted protein structures. Candidate sites are highlighted in orange, and protein domains are colored in pink and green. Labels begin with the amino acid position and residue in the reference sequence, followed by the amino acid substitution(s) identified in halophytes. Orthogroups with chi-square-tested, FDR-adjusted P-values < 0.05 and foreground 10 > ω > 1 were considered as PSGs and REGs in halophytes. Positively selected amino acid sites were inferred as sites with a Bayes empirical Bayes posterior probability > 0.95.

We next examined genes from halophytes containing sites with signatures of positive selection. Despite limited overlap in genes with positively selected sites, there are many key functions associated with salt responses shared between Brassicaceae and Poaceae (Fig. 4C; Tables S14 and S15). These in Brassicaceae included *MAPKKK20 (M3K20), PHL1, PIAL1*, cation exchanger *CAX5*, and aquaporin *TIP1;1*. Similarly identified genes in Poaceae prominently featured membrane transporters involved in ion and water homeostasis, including *NHX1, NHX6, KEA2, KEA5, KCO2, AVP,* and *TPC1*. Both families also showed positive selection in genes associated with stress-response hormonal signaling and oxidative stress mitigation. Although the specific genes differed between families, their functional associations point to recurrent evolutionary recruitment of related pathways during adaptation to saline environments. Among the genes that are fast-evolving and with positively selected sites targeted independently across diverse halophytes, RALF27 (Rapid Alkalinization Factor) in Brassicaceae and the NHX1 cation/H⁺ antiporter in Poaceae are notable because of their established roles in salt stress responses (*17*, *18*). RALF peptides participate in cell wall integrity signaling in regulating growth and salt tolerance, while NHX1 contributes to intracellular cation and pH homeostasis. We mapped recurrently selected amino acid sites identified across halophytes onto the corresponding Arabidopsis or rice reference proteins (Fig. 4D). Different halophyte lineages exhibited signatures of positive selection in distinct regions of the protein sequence, with some regions independently targeted by multiple halophytes (Fig. 4D; Table S16). We have identified genes that showed rapid evolutionary rates and recurrent signals of positive selection but are poorly characterized, making them promising candidates for uncovering novel functions repeatedly recruited during halophyte evolution (Table S13).

### Shared functions, distinct paths: convergent salt adaptation and divergent genomic evolution in halophytes compared to crop relatives

Modern crop breeding has prioritized yield under favorable conditions, often with trade-offs in stress resilience, whereas wild relatives retain genetic diversity and adaptive traits shaped by selection across diverse natural environments (*19–21*). Comparative analyses of crops and their halophytic relatives across multiple lineages can uncover convergent genomic signatures of salt adaptation and reveal underexplored targets for enhancing crop resilience (*3*, *22–24*). We selected six halophytes and their crop relatives spanning eudicots and monocots, including congeneric and conspecific pairs, to examine differences in gene family sizes, basal expression, and evolutionary rates associated with selection. These included *Beta vulgaris* subsp. *vulgaris* and *B. vulgaris* subsp. *maritima*, *Glycine max* and *G. soja*, *Hordeum vulgare* and *H. marinum*, *Oryza sativa* and *O. coarctata*, *Solanum lycopersicum* and *S. chilense*, and *Vigna radiata* and *V. marina*.

We compared orthogroup expansions and contractions to identify convergent patterns of gene family size evolution in the wild halophytes (Fig. 5A; data S9 and S10). Rapid gene family expansion beyond the expected rate for a lineage may indicate preferential retention of duplicated genes under selection, whereas contraction may reflect relaxed selection or adaptive gene loss, potentially including the loss of regulatory components that constrain responses to changing environments (*25*, *26*).

**Fig. 5.**
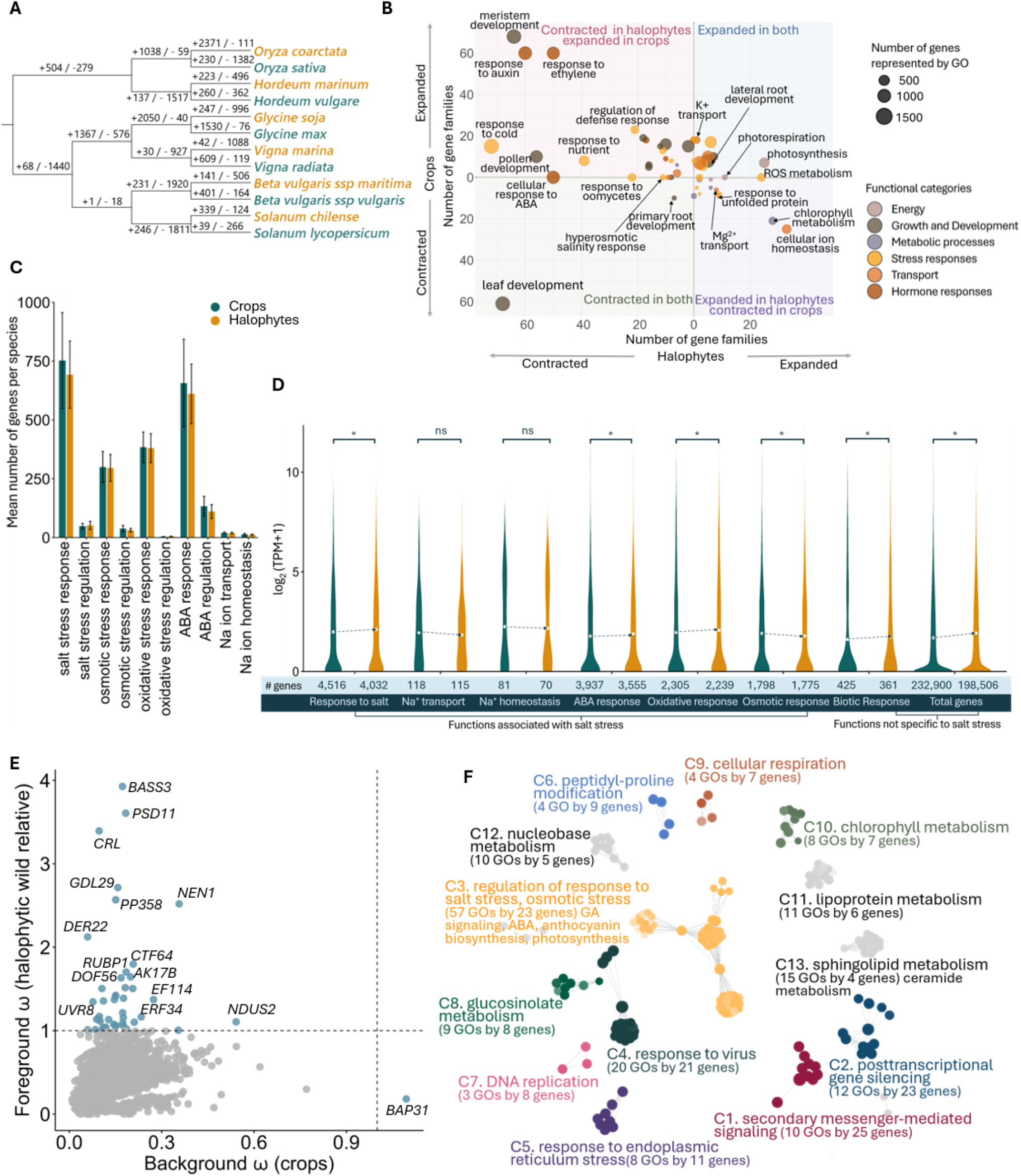
Divergent evolutionary signals in crops and their congeneric halophytic wild relatives. **(A)** Gene family expansions and contractions among crops and their wild relatives included in the analysis. Expansion (positive values) and contraction (negative values) for gene families are indicated on branches. Crops are shown in green and halophytic wild relatives in gold. **(B)** Functional enrichment of expanded and contracted gene families between congeneric pairs of crops and their wild relatives. Each bubble represents an enriched biological function, and bubble size indicates the number of genes associated with each Gene Ontology (GO) term. The four quadrants distinguish functions expanded or contracted in crops and/or wild relatives. Colors indicate broad functional groups. **(C)** Mean number of genes per species in functional groups associated with salt responses. Halophytes (gold) and crops (green). Differences between groups were assessed using the Wilcoxon rank-sum test, with significance defined as Benjamini– Hochberg false discovery rate (FDR)-adjusted P < 0.05. Error bars indicate mean ± SD. **(D)** Basal expression distributions of genes assigned to salt-response functions and biotic response for halophytes (gold) and crops (green). Dashed arrows connect the mean basal expression and indicate the directional differences. Significant differences between halophytes and crops were tested using the Wilcoxon rank-sum test. Asterisks indicate Benjamini–Hochberg FDR-adjusted P-values < 0.05. ns: not significant. **(E)** Comparison of evolutionary rates (ω) estimated using a branch model, with halophytic wild relatives designated as the foreground and crops as the background. Orthogroups for which the alternative model provided a significantly better fit than the null model after chi-square testing and FDR correction (adjusted P < 0.05) are shown. Fast-evolving genes (ω > 1) are highlighted in blue. **(F)** Functional enrichment network of fast-evolving genes in halophytes relative to their crop relatives. Clusters represent broader functional categories, nodes represent GO terms, and labels identify representative enriched biological processes. Node size is proportional to gene number, and edges indicate shared genes between GO terms. All displayed clusters have FDR-adjusted P < 0.05. Darker node colors indicate greater enrichment significance.

Our analysis included 5,950 orthogroups represented in at least 50% of sampled species. Halophytes showed a net contraction of 3,321 orthogroups, compared with 2,329 in crops (Fig. 5A and Table S17). Functional enrichment revealed contrasting patterns in salt response-related gene families between the two groups (Fig. 5B and Table S18). Halophytes showed contractions in hyperosmotic salinity response and ABA, auxin, and ethylene signaling, but expansions in ROS metabolism and cellular ion homeostasis. In crops, ion homeostasis orthogroups instead contracted, while ABA, auxin, and ethylene response families expanded. These shifts align with the lower representation of genes across previously identified functional categories in halophytes (Fig. 5C and Table S19).

Halophytes also exhibited significantly higher basal expression than their crop relatives across multiple stress response functions, including salt response, ABA response, and oxidative stress response (Fig. 5D and Table S20), whereas Na⁺ transport and homeostasis showed no significant differences between the two groups. The elevated basal expression of genes associated with biotic stress further suggests that the differences between wild halophytes and crops extend beyond salt adaptation. Together, these patterns are consistent with crop domestication having broadly altered stress response expression, while wild relatives retain higher basal expression across diverse stress response pathways (*27*, *28*).

Because crop domestication can shape signatures of selection and evolutionary rates (*29*) we compared evolutionary rates between halophytes and their crop relatives in the selected set. Among 2,852 genes with significantly different evolutionary rates in crop relatives, 333 genes from 45 orthogroups showed evidence of positive selection (ω > 1; Fig. 5E and Table S21). Functional enrichment revealed a strong association with salt response pathways (Fig. 5F and Table S22). Many genes captured in the functional cluster C3 included responses to salt and osmotic stress, ABA biosynthesis, and regulation of anthocyanin biosynthesis, including transcription factors of stress regulation. These findings contrast with Brassicaceae and Poaceae halophyte comparisons (Fig. 4B), in which salt-associated functions were enriched among slow-evolving genes in halophytes. Across the four families represented in the crop–halophyte pairs, however, salt-associated functions were enriched among fast-evolving genes in halophytes. This contrast suggests that salt adaptation involves both conserved mechanisms within lineages (Fig. 4B) and fast-evolving genes across lineages (Fig. 5E and F). Such differences may reflect the distinct evolutionary routes through which halophytes achieve salt tolerance.

## Discussion

Comparative genomic studies have associated salt adaptation with gene family expansion, duplication, and accelerated molecular evolution. For example, analyses using *Puccinellia tenuiflora* have identified expanded and positively selected K⁺ uptake gene families, while genome studies of *Suaeda salsa* and *Spartina alterniflora* have reported expansions of genes associated with ion transport and salt responses (*30–32*). Additionally, studies of *Tamarix chinensis* and *Schrenkiella parvula* have linked salt adaptation to gene duplication in salt transport and response pathways (*33*, *34*). Despite these lineage-specific findings, our broad analysis across angiosperm families reveals a contrasting pattern: halophytes generally possess fewer gene copies in gene families associated with salt stress shared across species, while these genes exhibit higher and more tightly constrained basal expression and comparatively conserved evolutionary rates (Fig. 2-4). This contrast suggests that salt adaptation need not depend on pervasive gene family expansion or accelerated protein evolution. Our study presents a ubiquitous alternative genomic strategy for maintaining salt-response capacity independently in diverse lineages. Importantly, lineage-specific expansions and positive selection in specific genes remain compatible with this broader pattern, indicating that adaptive innovation in individual pathways can occur within an overall genomic context of reduced redundancy and evolutionary constraint.

Our analysis often included genes belonging to large orthogroups shared across deep evolutionary nodes, capturing broadly conserved components of salt response networks rather than lineage-specific gene acquisitions. These genes may function within regulatory networks, in which coordinated activity and balanced gene dosage are important for maintaining cellular homeostasis. Because these orthogroups span deep evolutionary divergences, their conservation may reflect shared genomic foundations of salt adaptation across angiosperms, whereas lineage-specific expansions and fast-evolving genes may represent more recent or specialized adaptations. Our categorization of halophytes clusters species with distinct physiological strategies, including salt exclusion, salt accumulation, and salt secretion through specialized salt glands (*8*, *35*, *36*) into a single foreground (Fig. 1). Thus, the convergent signatures identified in this study reflect patterns detectable across diverse physiological strategies. Further analyses of specific phenotypic subgroups may reveal genomic signatures associated with distinct mechanisms of Na⁺ transport and ion homeostasis that contribute to the halophyte–non-halophyte discrimination observed in Figs. 2–5. Together, these findings suggest that convergent halophytism can involve both the long-term retention and regulation of shared genetic components and lineage-specific genomic innovations, with their relative contributions depending on evolutionary scale and functional context.

Our analyses are subject to several limitations inherent to comparative genomics across highly diverged plant lineages. Differences in genome assembly quality, gene annotation, and orthogroup inference may affect gene copy number estimates, while alignment errors and uncertain homology assignments can influence estimates of evolutionary rates and positive selection. Because our evolutionary analyses rely primarily on substitution patterns in coding sequences, they may not capture selection acting through gene regulation, gene dosage, structural variation, or other mechanisms independent of sequence substitution. Moreover, genes grouped into the same orthogroup may have diverged functionally, particularly where functional characterization remains limited in non-model species. Although we accounted for coding potential, some annotated copies may represent pseudogenes or nonfunctional remnants, while highly diverged or lineage-specific genes may have been missed or excluded from orthogroups, potentially obscuring clade-specific expansions and adaptive innovations. Although individual errors in genome assembly, gene annotation, orthogroup assignment, or evolutionary inference may affect specific genes or lineages, their influence on broad patterns observed across this diverse large-scale dataset of >200 genomes with similar representation of halophytes and non-halophytes may be limited. Nevertheless, these potential sources of error warrant caution when interpreting gene copy representation and evolutionary constraint as direct proxies for functional contributions to salt tolerance.

Our curated collection of orthogroups and associated evolutionary and expression profiles provides a valuable resource for identifying candidate genes and conserved gene networks underlying salt tolerance. Our findings reveal recurrent genomic trends shared across phenotypically diverse halophytes defined by a common salinity tolerance threshold, providing a genomic foundation for prioritizing candidates whose contributions to specific salt-tolerance strategies await functional dissection. In particular, gene families conserved across deep evolutionary nodes, shared among many lineages, and associated with multiple stress response mechanisms offer promising candidates for transferring stress tolerance traits from wild halophyte relatives to crops.

## Materials and Methods

### Data collection and curation

Halophytes were defined as plant species reported in the literature to survive at NaCl concentrations greater than 200 mM. Halophyte subtypes (salt accumulators, excluders, recretohalophytes, and succulent dilutors) could not be consistently annotated across species because many non-model species in our catalog lack systematic physiological characterization. We therefore used growth at ≥200 mM NaCl as a unified physiological trait encompassing diverse salt-tolerance strategies (*1*, *37*). Publicly available coding sequences (CDS) for halophytes and non-halophytes were retrieved from genome repositories, including NCBI (https://www.ncbi.nlm.nih.gov/), Phytozome-JGI (https://phytozome-next.jgi.doe.gov/), NGDC (https://ngdc.cncb.ac.cn/gwh), and published literature. For species without publicly available CDS files, genome assemblies were downloaded and used for gene model prediction. Genome completeness was assessed using BUSCO v5.2.2 with the embryophyta database (*38*). Genomes with BUSCO completeness scores lower than 80% were discarded. Genomes were repeat-masked using RepeatMasker v4.1.5 (*39*) and gene models were predicted from soft-masked draft genomes using BRAKER 3.0.8 with RNA-seq evidence (*40*).

CDS datasets with BUSCO completeness scores greater than 80% were retained for further curation. The coding potential of predicted gene models was evaluated using CPC2 (coding probability > 50%) (*41*). For each species, gene models that passed the CPC2 coding-potential threshold were renamed using a consistent gene ID format to facilitate downstream analyses. A second round of BUSCO assessment was performed after CPC2 filtering, and CDS datasets with BUSCO completeness scores greater than 80% were retained as the final dataset. The final species list contains 232 angiosperm species, including 114 halophytes and 118 non-halophytes.

Raw transcriptome data were collected from the NCBI SRA database. For each species, non-stress/control shoot tissue samples representing basal expression levels were selected when available. Shoot tissue was selected because non-stress shoot RNA-seq was substantially more available across the 212 species than non-stress root RNA-seq, enabling balanced sampling of 107 halophytes and 105 non-halophytes. Shoot regulation of ion homeostasis is itself central to salt tolerance because toxic Na⁺ accumulation in photosynthetic tissue drives much of the cellular damage caused by salinity. Transcriptomes generated using BGISEQ and Illumina platforms were included. When available, datasets with read lengths greater than 100 bp and at least around 30 million reads per species were selected.

### Gene Model Functional Classification

TransDecoder v5.5 (https://github.com/TransDecoder/TransDecoder) was used to predict open reading frames and corresponding protein sequences from the curated CDS datasets. Those were subsequently annotated using Trinotate v3.2.2 (*42*), including sequence-similarity searches against BLAST databases and protein-domain identification using the Pfam database. When applicable, SignalP was used to identify putative signal peptides. Gene models were additionally annotated using Gene Ontology (GO) and KEGG databases to assign functional annotations and pathway associations.

### Orthology inference and species tree construction

Orthogroups were inferred using OrthoFinder v3.0.1b1 (*43*) separately for the angiosperm clade, Brassicaceae species, Poaceae species, and crop species with their wild relatives. Orthogroup inference was performed using both CDS and protein datasets for each group using default parameters (multiple sequence alignments: MAFFT v.7.525, phylogenetic tree inference: FastTree v.2.2.0). The species tree obtained for the angiosperm clade from OrthoFinder was manually curated to be consistent with the Kew Gardens taxonomy tree (*44*). Orthogroups were functionally annotated using a custom Python v3.9 (https://www.python.org/downloads/release/python-390/) script that integrated the functional annotations of genes assigned to each orthogroup across species.

### Orthology comparison between halophytes and non-halophytes

For each species, OrthoFinder-derived gene counts were summarized into four orthology categories: paralogs shared among species, single-copy orthologs, species-specific paralogs, and species-specific single-copy genes. The count for each category was normalized as a percent of the total number of genes of each species. Mean percent was compared between halophytes and non-halophytes for each category using the Wilcoxon rank-sum test, followed by Benjamini– Hochberg false discovery rate correction at FDR-adjusted P-values < 0.05.

Species clustering was performed using orthogroups represented in ≥25% or ≥50% of species to mitigate sparsity across 18 plant orders. UMAP analysis was performed using the Python v3.9 package umap-learn (*45*) with the following parameters: random_state = 42, n_neighbors = 10, and min_dist = 0.6.

We obtained custom images from iNaturalist (https://www.inaturalist.org/) for selected halophytes and trained ChatGPT (https://chatgpt.com/) to create plant illustrations. All generated illustrations were manually reviewed. These illustrations were used only for visual representation and were not included in downstream quantitative analyses.

### Gene counts for functions associated with salt

Custom Python scripts (v3.9) were used to identify and count genes associated with selected biological functions based on Gene Ontology (GO) identifiers. For each species, the number of genes assigned to each function was counted. Differences in mean gene number per species between halophytes and non-halophytes were tested using a Wilcoxon rank-sum test, followed by Benjamini–Hochberg false discovery rate correction at FDR-adjusted P-values < 0.05.

To compare relative gene allocation between halophytes and non-halophytes in Fig. 2, D and E, for each species, gene counts for each function were normalized by the total number of annotated genes. This normalized value was calculated as:

*number of genes assigned to a function / total number of annotated genes per species*

For each plant family containing at least two halophyte and two non-halophyte species, the mean normalized gene allocation was calculated separately for halophytes and non-halophytes. The difference in mean gene allocation was then calculated as:

*Mean(normalized gene allocation)_halophytes_ − Mean(normalized gene allocation)_non-halophytes_*

Positive values indicate higher relative gene allocation in halophytes, whereas negative values indicate higher relative gene allocation in non-halophytes. Gene allocation differences were compared between functions associated with salt stress and functions not specific to salt stress using Student’s t-test, with statistical significance set at P < 0.05.

### Separation of halophytes and non-halophytes by PCA

Principal component analysis (PCA) was performed in R v4.5.0 (*46*) using the prcomp function (*47*) PCA was performed using gene counts for selected biological functions and orthogroup-level gene-count matrices. Separation between halophytes and non-halophytes along individual principal components was evaluated using the Wilcoxon rank-sum test and Cohen’s d. To identify the pair of principal components that best separated halophytes and non-halophytes, binomial logistic regression models were fitted using PC scores, and model performance was evaluated using the area under the receiver operating characteristic curve (AUC) with the pROC package (*48*).

PCA was performed using four datasets: (1) gene counts for selected salt-associated functions for each species, (2) gene counts of orthogroups with greater than 50% species representation, (3) basal level expression of salt-associated genes in each species, and (4) basal level expression of orthogroups with greater than 50% species representation. For the orthogroup-based expression PCA, the highest basal expression value among genes within each orthogroup was selected for each species.

The contribution of each function or orthogroup to group separation was estimated from the PCA loading matrix. For each function or orthogroup, combined loading strength was calculated using the two principal components that best separated halophytes and non-halophytes: √(loading_PCx^2^ + loading_PCy^2^), where loading_PCx and loading_PCy represent the loadings of a function or orthogroup on the selected principal components. Functions and orthogroups with the highest combined loading strengths were considered the major contributors to halophyte/non-halophyte separation.

### Basal level expression inference

RNA-seq data generated under control conditions were mapped against reference gene models using Salmon v.1.5.1 (*49*). Expression levels were normalized as transcripts per million (TPM). To evaluate the relationship between gene number and basal expression, the average log_2_(TPM + 1) value was calculated for each species and compared against the total number of genes. Linear regression was used to assess the relationship between total gene number and average basal expression separately for halophytes and non-halophytes. The coefficient of variation was calculated as the standard deviation divided by the mean expression per orthogroup, separately for halophytes and non-halophytes. The mean coefficient of variation was compared between halophytes and non-halophytes using the Wilcoxon rank-sum test, with statistical significance set at P < 0.05.

For each selected biological function, basal expression values of genes assigned to that function were extracted for each species. Mean expression differences between halophytes and non-halophytes were tested using the Wilcoxon rank-sum test, adjusted for multiple testing using false discovery rate correction. Statistical significance was set at Benjamini–Hochberg FDR-adjusted P-values < 0.05. The coefficient of variation of the basal level expression for the selected functions was also calculated for halophytes and non-halophytes.

### Relative evolutionary rates and positive selection

Evolutionary rates were estimated for orthogroups from Brassicaceae and Poaceae. Orthogroups with greater than 50% species representation within each family were used for the analysis. CODEML in the PAML v4.10.7 package (*50*), which estimates the ratio of nonsynonymous to synonymous substitution rates (ω = dN/dS), was used to compare evolutionary rates between halophytes and non-halophytes. Branch models and branch-site models were used to identify fast- and slow-evolving genes and positively selected genes (PSGs), respectively. For both analyses, halophyte branches were assigned as foreground branches, whereas non-halophyte branches were assigned as background branches.

Protein sequences from orthogroups were aligned using MAFFT v.7.525 (*51*) and the alignments were converted into their corresponding codon alignments using their respective CDS orthogroup as a reference in PAL2NAL v.14 (*52*). Codon alignments were converted to PHYLIP format using FASTAtoPHYL.pl for CODEML analysis. Gene trees were further processed using custom Python scripts to label foreground and background branches.

For the branch model analysis to test fast- and slow-evolving genes, a one-ratio null model, in which all branches share the same ω value, was compared with a two-ratio alternative model, in which foreground and background branches were allowed to have different ω values. The null model was run with model = 0, fix_omega = 0, and omega = 1.5, whereas the alternative model was run with model = 2, fix_omega = 0, and omega = 1.5. Likelihood ratio tests were used to compare the null and alternative models using LRT = 2 × (lnLalternative − lnLnull). P-values were calculated using a chi-squared test and adjusted for multiple testing using false discovery rate correction. Orthogroups with FDR-adjusted P-values < 0.05 were considered as orthogroups best fit to the two-ratio model. The orthogroups with abnormally high ω values > 10 were discarded (*53*).

Genes were ranked according to foreground ω values estimated by the branch model. Knee-point analysis was used to identify inflection-point thresholds for selecting orthogroups with the fastest and slowest evolutionary rates in halophyte foreground branches. Orthogroups above the upper knee-point threshold and foreground ω > background ω were considered candidate genes from the fast-evolving, whereas orthogroups below the lower knee-point threshold were considered slowly evolving or conserved candidate genes.

For positive selection analysis, the branch-site model was used to identify genes with evidence of positive selection on foreground branches. The null model was run with model = 2, NSsites = 2, fix_omega = 1, and omega = 1, whereas the alternative model was run with model = 2, NSsites = 2, fix_omega = 0, and omega = 1.5. The best-fitting model was assessed using likelihood ratio tests, and P-values were calculated using the chi-squared test and adjusted using false discovery rate correction. Orthogroups with FDR-adjusted P-values < 0.05 and foreground 10 > ω > 1 were considered as positively selected genes in halophytes. Positively selected amino acid sites were inferred using the Bayes empirical Bayes method (BEB), and sites with BEB posterior probability > 0.95 were considered candidate sites under positive selection.

### Mapping of positively selected sites on protein models

AlphaFold protein structures were downloaded from the AlphaFold database (https://alphafold.ebi.ac.uk/). The amino acid structure annotation was done using PyMOL v3.0.0 (https://www.pymol.org/)

### Expanded and contracted gene families

Orthogroups with greater than 50% species representation were used to identify expanded and contracted gene families using CAFE v.5.1.0 (*54*) with default parameters. For each orthogroup, expansion and contraction events were summarized separately for halophyte and non-halophyte lineages. Orthogroups with more expansion events than contraction events in halophyte lineages were classified as expanded in halophytes. Orthogroups with more contraction events than expansion events in halophyte lineages were classified as contracted in halophytes. The same approach was applied to non-halophyte lineages.

### Functional enrichment analysis

Functional enrichment analysis for expanded gene families, contracted gene families, REG, and PSG was performed using BiNGO (*55*) implemented in Cytoscape (*56*) and GOMCL (*57*) using custom-made gene association files for each plant family orthogroup set as a background. For each plant family, orthogroup-based gene-association files were generated using a custom Python script and used as the background. Enrichment was tested at adjusted p-values < 0.05 with FDR correction applied.

## Acknowledgments

We thank Drs. John Larkin, Jeremy Brown, Sarah Mathews, Guannan Wang, Pramod Pantha, and lab members Thu Nguyen, Richard Garcia, Isabel Carmen Johnson, Chamara Mendis, and Mohomed Elgallad for their constructive feedback. The authors also acknowledge the LSU High Performance Computing services for providing computational resources needed for data analyses. ChatGPT (OpenAI) was used to create plant vector illustrations in Fig. 1A. This work was shaped in part by discussions with Dr. John M. Cheeseman (1948–2026) and by the body of research he helped advance. MD gratefully acknowledges his lasting scientific influence and the important legacy of his contributions to the field. This work was supported by the US National Science Foundation (EDGE 1923589) and the Department of Energy (BER DE-SC0022985) awards.

## Author contributions

Conceptualization: MD

Methodology: SW, MD

Investigation: SW

Visualization: SW

Funding acquisition: MD

Project administration: MD

Supervision: MD

Writing – original draft: SW

Writing – review & editing: SW, MD

---

**Fig. S1.**
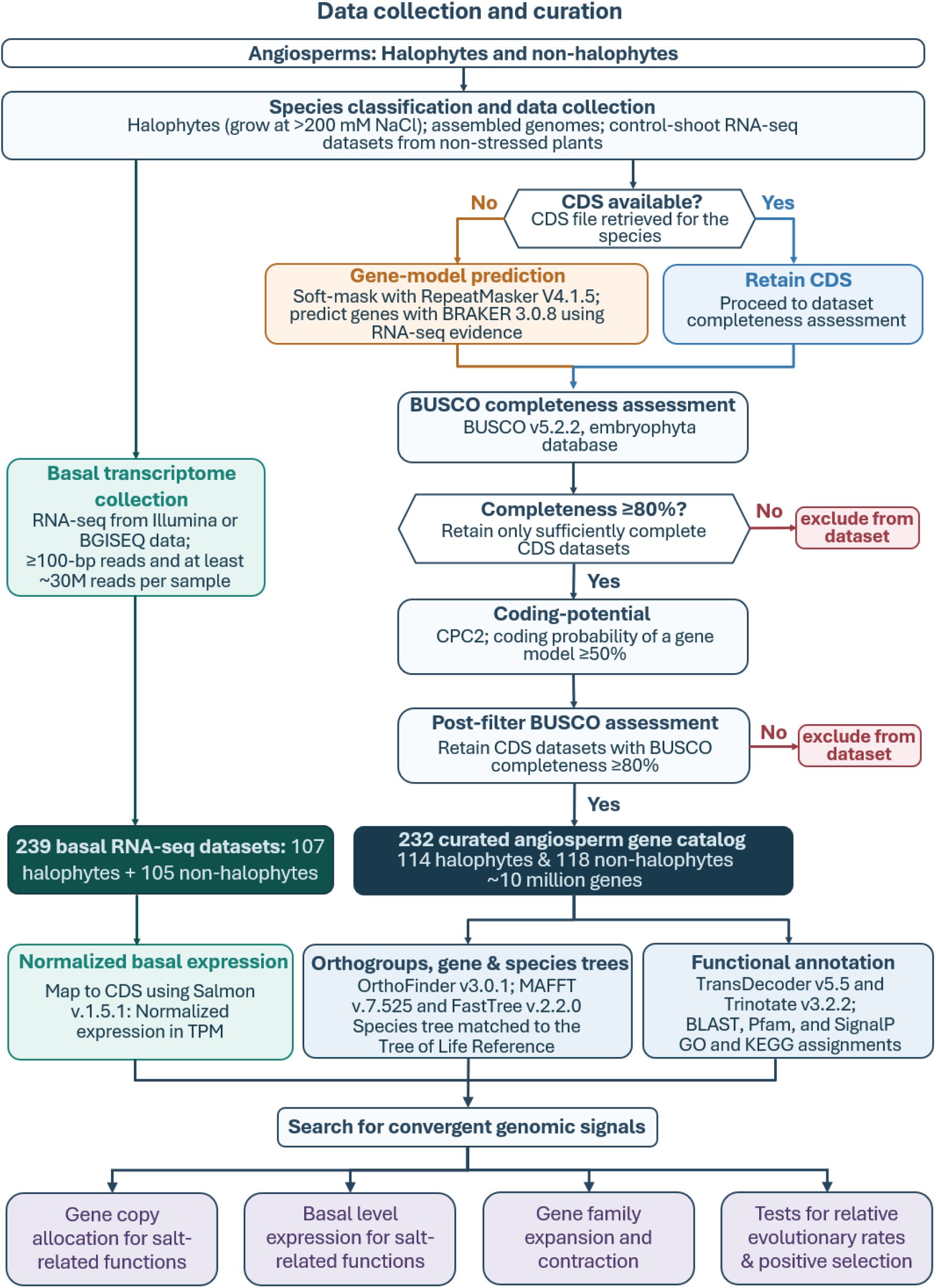
Summary workflow to construct the catalog of angiosperm protein-coding genes.

**Fig. S2.**
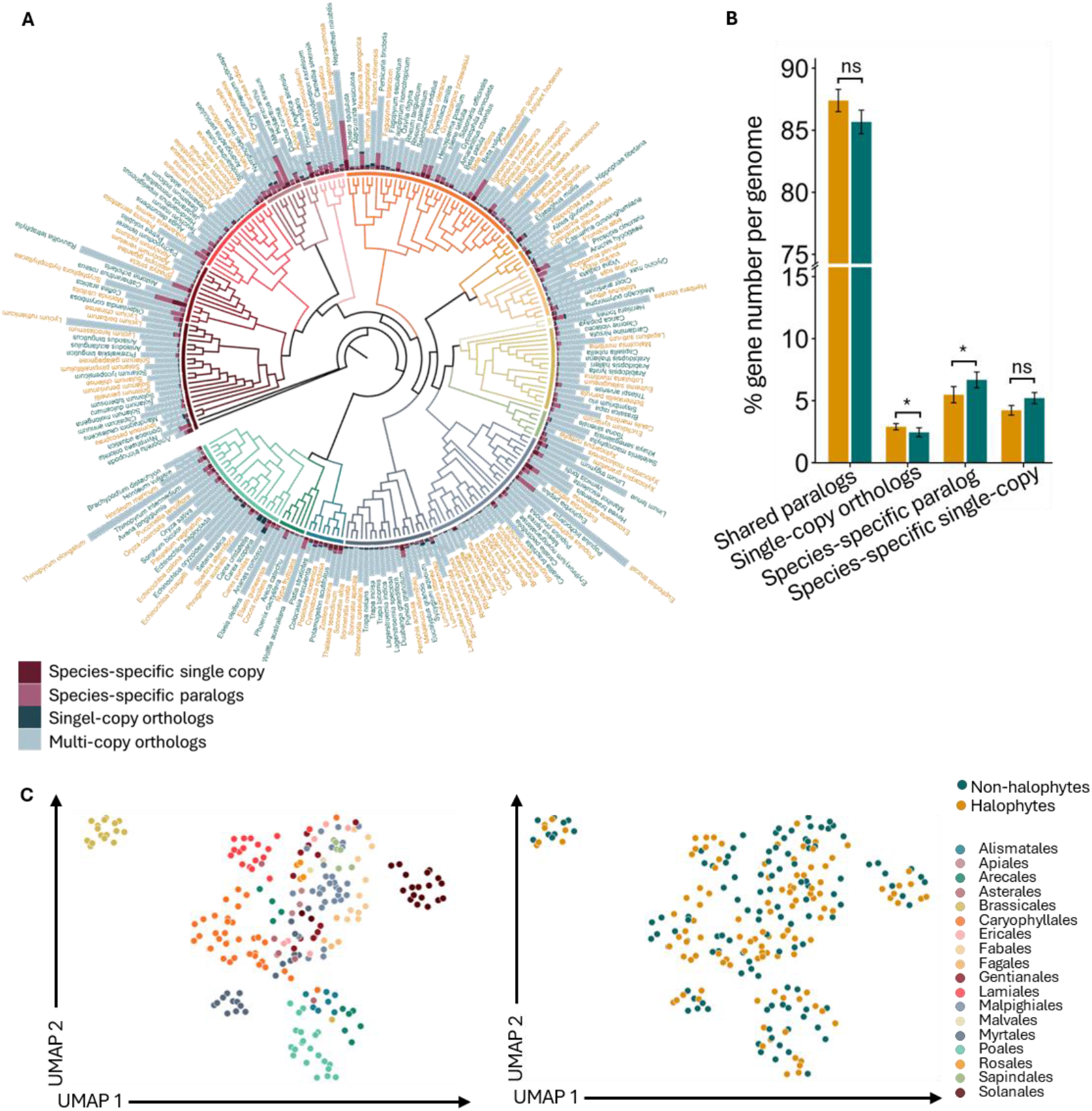
Gene composition and phylogenetic relationships among the halophytes and non-halophytes. **(A)** The species tree shows the relationship of angiosperm genomes studied. Bars around the phylogeny represent the number of genes assigned to ortholog classes in each species. **(B)** Percentage of genes assigned to ortholog classes in each genome. Total number of genes: 150,173 in shared paralogs, 68,703 in single-copy orthologs, 108,155 in species-specific paralogs, and 418,123 in species-specific single-copy (which were excluded from orthogroup assessments). Asterisks indicate significant differences at FDR-adjusted P < 0.05 based on Wilcoxon rank-sum tests with Benjamini–Hochberg correction; ns, not significant. Error bars indicate mean ± SD. **(C)** Clustering of species based on the number of genes in orthogroups represented in >50% of species. The two panels are colored by plant order and halophytic status, respectively, showing that species cluster primarily according to phylogenetic relationships.

**Fig. S3.**
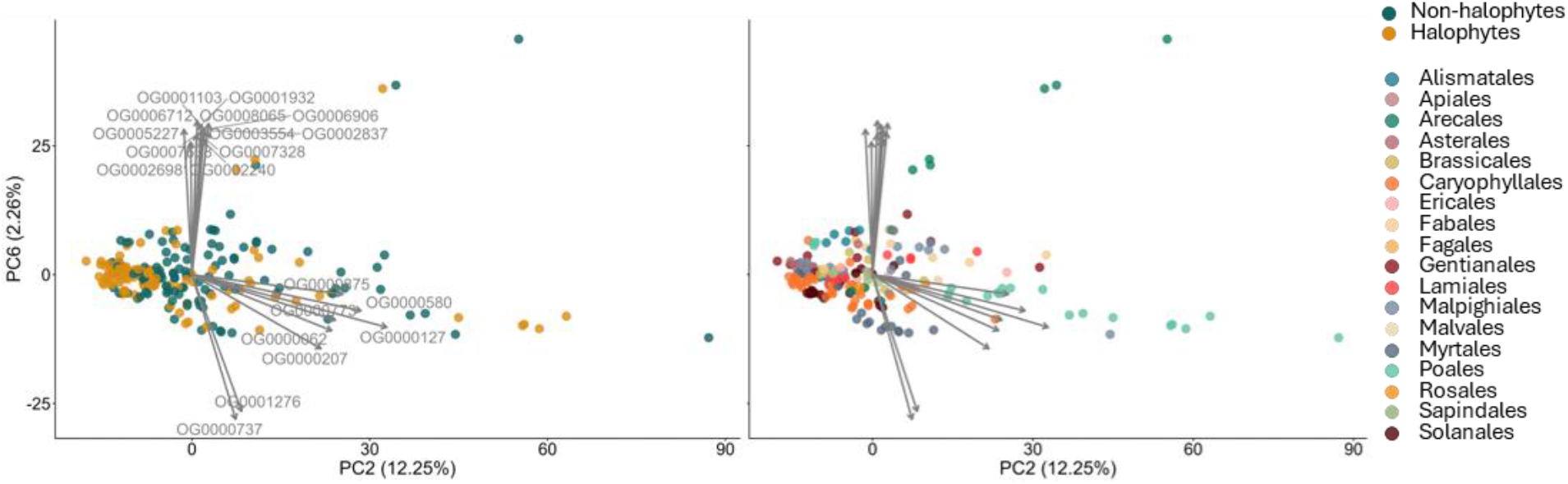
Gene composition in common orthogroups separates halophytes from non-halophytes. Principal components (PCs) most influential in separating halophytes from non-halophytes based on gene composition in orthogroups represented in >50% of sampled species. PCs selected based on Cohen’s d and Benjamini–Hochberg false discovery rates. Species are colored by halophytic status (left) or plant order (right). Arrows indicate the 15 orthogroups contributing most to the separation of species, with arrow length proportional to the magnitude of each orthogroup’s contribution.

**Fig. S4.**
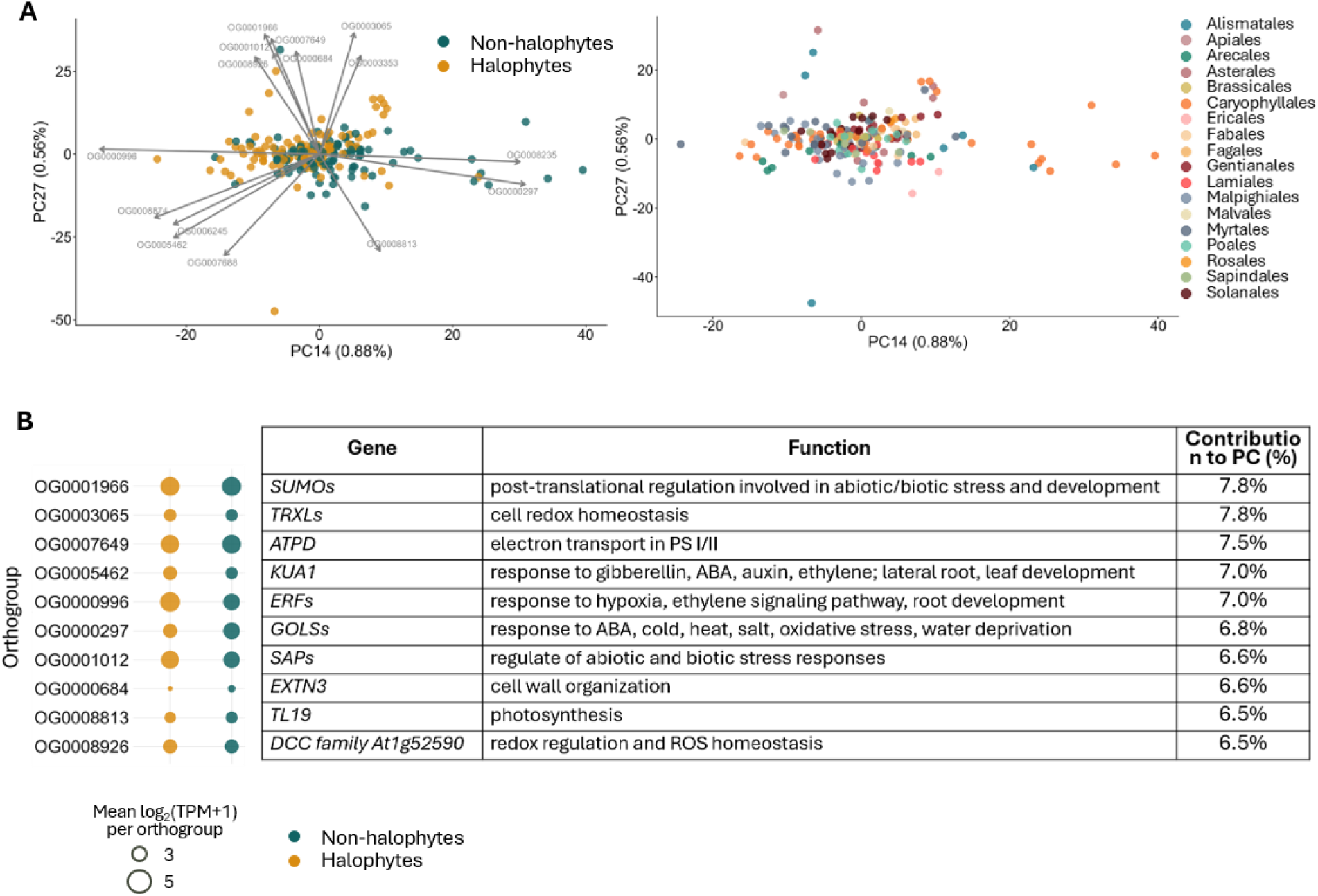
Basal expression in common orthogroups separates halophytes from non-halophytes. **(A)** Principal components (PCs) most influential in separating halophytes from non-halophytes based on basal expression of genes in orthogroups which represented in >50% of sampled species. PCs selected based on Cohen’s d and Benjamini–Hochberg false discovery rates. Species are colored by halophytic status (left) or plant order (right). Arrows indicate the 15 orthogroups contributing most to the separation of species, with arrow length proportional to the magnitude of each orthogroup’s contribution. **(B)** Top ten orthogroups contributing most to the separation of halophytes and non-halophytes in (A), with orthogroup IDs, functional annotations, and percentage contributions to the PCs. Bubble sizes indicate the mean log_2_(TPM+1) per orthogroups.

**Fig. S5.**
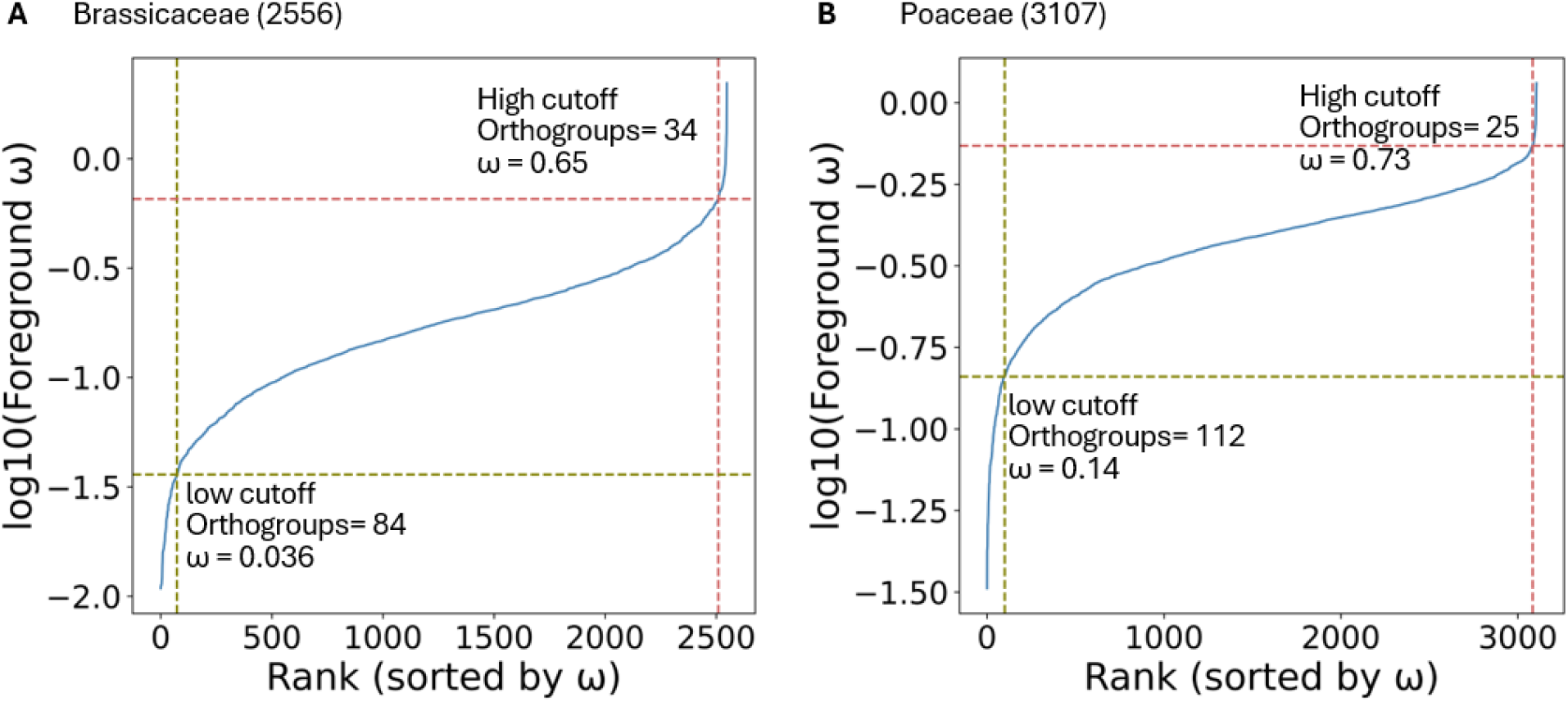
Cumulative contribution of ω values ranked to select genes with the fastest and slowest evolutionary rates in Brassicaceae and Poaceae orthogroups with a 50% species representation. **(A)** Brassicaceae and **(B)** Poaceae. Upper and lower inflection points were used as cutoffs for fast- and slow-evolving genes, respectively.

